# An Optimised GAS-pharyngeal cell biofilm model

**DOI:** 10.1101/2020.11.25.399196

**Authors:** Heema K. N. Vyas, Jason D. McArthur, Martina L. Sanderson-Smith

## Abstract

Group A *Streptococcus* (GAS) causes 700 million infections and accounts for half a million deaths per year. Biofilm formation has been implicated in both pharyngeal and dermal GAS infections. *In vitro*, plate-based assays have shown that several GAS M-types form biofilms, and multiple GAS virulence factors have been linked to biofilm formation. Although the contributions of these plate-based studies have been valuable, most have failed to mimic the host environment, with many studies utilising abiotic surfaces. GAS is a human specific pathogen, and colonisation and subsequent biofilm formation is likely facilitated by distinct interactions with host tissue surfaces. As such, a host cell-GAS model has been optimised to support and grow GAS biofilms of a variety of GAS M-types. Improvements and adjustments to the crystal violet biofilm biomass assay have also been tailored to reproducibly detect delicate GAS biofilms. We propose 72 h as an optimal growth period for yielding detectable biofilm biomass. GAS biofilms formed are robust and durable, and can be reproducibly assessed via staining/washing intensive assays such as crystal violet with the aid of methanol fixation prior to staining. Lastly, SEM imaging of GAS biofilms formed by this model are resemblant of those previously found on excised tonsils of patients suffering chronic pharyngo-tonsillitis. Taken together, we outline an efficacious GAS biofilm pharyngeal cell model that can support long-term GAS biofilm formation, with biofilms formed closely resembling those seen *in vivo*.

## Introduction

*Streptococcus pyogenes* (Group A *Streptococcus*; Group A *streptococci*; GAS) is a Gram-positive human pathogen known to cause an array of infections ranging from mild infections of the skin and throat, to more serious and life threatening conditions such as necrotising fasciitis and numerous autoimmune sequelae [1]. GAS infections are a considerable burden on global healthcare systems with high rates of patient mortality and morbidity [2].

GAS has been found to form biofilms in the tonsillar crypts of patients with GAS pharyngitis and in the skin lesions of GAS impetigo sufferers [3,4]. *In vitro*, it has been demonstrated that GAS biofilm formation is strain dependent. And among isolates of the same serotype, biofilm forming capacities are oftentimes found to differ considerably [5]. As highlighted in Table 1, *in vitro* plate-based studies have implicated several GAS virulence factors (M protein, capsule, pili, SpeB, CovS, and quorum sensing peptides) in biofilm formation [6]. These findings have contributed substantially to our current understanding of GAS biofilms and their involvement in GAS pathogenesis and disease. However, much of this work has been conducted on abiotic surfaces (plastic, glass, and silicone). To date, few studies have used host matrix components like collagen, fibronectin, or fibrinogen as surface coatings for GAS biofilm studies [7–10]. Moreover, there is currently no methodology or protocol widely recognised as the gold standard for GAS biofilm formation. The variability among methods, and limited use of host factors in *in vitro* plate-based GAS biofilm models found in previous studies has been summarised in Table 1. This table highlights that only three *in vitro* plate-based studies utilising epithelial monolayers to grow and support GAS biofilms have been published.

**Table 1.**
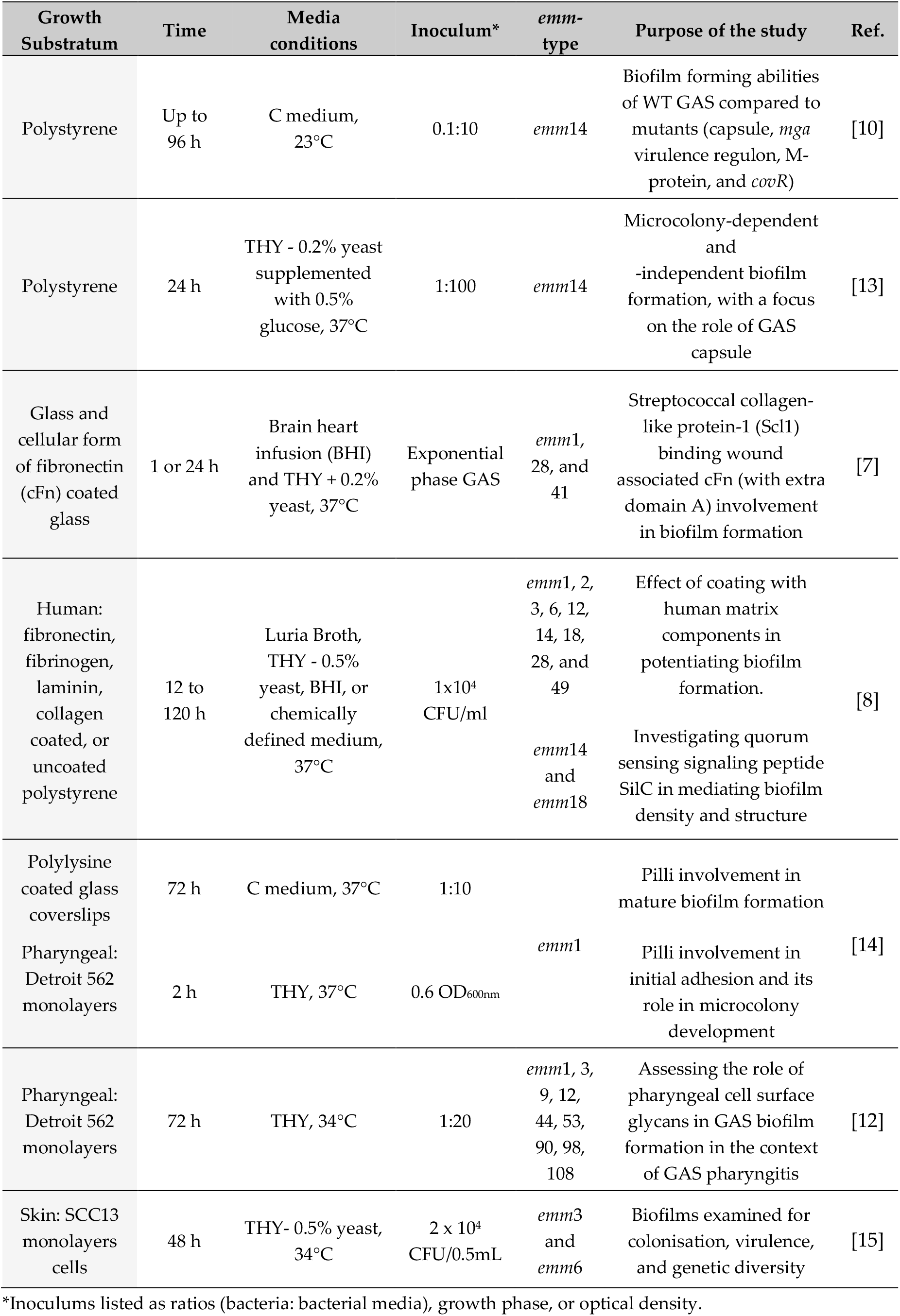
Examples of *in vitro* plate-based models used for the study of GAS biofilm formation.

There is a need for GAS biofilm models to better incorporate host factors, as it has been found that failure to do so has significant effects on the biofilms formed, with the overall arrangement and architecture of biofilms and their virulence gene expression, found to noticeably differ from that of biofilms formed *in vivo* [10]. Moreover, tissue tropism displayed by differing GAS isolates towards the throat and skin, which are vastly different epithelial landscapes and environments [11], will influence GAS adherence, colonisation, and subsequent biofilm formation. Thus, the incorporation of relevant host epithelial substratum, and an overall mimicking of the host environment should be a consideration in GAS biofilm modelling.

Here, a GAS-pharyngeal cell biofilm model has been optimised to cultivate robust GAS biofilms that can be used for a diverse set of GAS M-types. We also present optimised steps and tips for increased biofilm integrity and reproducibility when performing staining assays like crystal violet. This model has since been used to effectively assess the role pharyngeal cell surface glycans play in GAS biofilm formation and clearly demonstrates the importance of mimicking the epithelial environment in these studies [12].

## Materials and Methods

### GAS and culture conditions

GAS strains used in this study (Table 2) are clinical GAS isolates, with each strain representative of a discrete GAS *emm*-type [16–18]. GAS was grown on horse blood agar (HBA) plates (Oxoid, UK) or Todd Hewitt agar supplemented with 1% (w/v) yeast (THYA) (Difco, Australia). Static cultures of GAS were grown overnight in Todd Hewitt broth supplemented with 1 % (w/v) yeast (THY). GAS was cultured, maintained, and biofilms formed at 34°C to mimic conditions more closely seen in the *in vivo* pharyngeal environment as described by Marks, *et al.* [19].

**Table 2.** GAS strains utilised in this study, their *emm*-types, and clinical source.

| M-type | Strain | Clinical source | Ref. |
| --- | --- | --- | --- |
| M1 | 5448 | Invasive infection: Necrotising fasciitis and toxic shock | [18] |
| M12 | PRS-8 | Superficial infection: persistent Pharyngeal pus/sinusitis | [17] |
| M3 | 90254 | Invasive infection | [17] |
| M98 | NS88.2 | Invasive infection: Blood (bacteraemia) | [16] |
| M108 | NS50.1 | Superficial infection: Wound | [16] |

### Human pharyngeal cell culture conditions

Detroit 562, a human pharyngeal epithelial cell line (CellBank Australia, Australia), was cultured in Dulbecco’s Modified Eagle Medium (DMEM) F12 (Invitrogen, Australia), supplemented with 2 mM L-glutamine (Gibco, Life Technologies, UK) and 10 % (v/v) heat inactivated foetal bovine serum (FBS) (Bovogen Biologicals, Australia) in cell culture flasks at 37°C, 5% CO_2_ – 20% O_2_ atmosphere.

### Pharyngeal cell monolayer formation

Detroit 562 pharyngeal cell monolayers form the substratum for GAS biofilm growth. An outline of the process and the monolayers formed is depicted in Fig. 1.

- The wells of a 96-well flat bottom cell culture microtiter plate (Greiner Bio-One, Germany) were coated with 50 μL of 300 μg/mL Collagen I from rat tail (Gibco, Life Technologies, UK) prepared in pre-chilled, sterile 17.4 mM acetic acid solution. The plate was incubated for 1 h, 37°C, 5% CO_2_ – 20% O_2_ atmosphere.
- After 1 h, excess collagen was removed, and the wells seeded with 150 μL Detroit 562 cell suspension (2×10^5^ cells/mL) and cultured for 48 h (to achieve ~95% confluency).
- Monolayers were washed once with 200 μL of sterile PBS, and fixed with 50 μL sterile 3.7% paraformaldehyde (PFA) (w/v) for 20 min.
- Once cells were fixed, PFA was removed and wells washed twice with 200 μL of PBS.
- Monolayers can be used immediately, or stored at 2-8°C for up to two weeks (with monolayers kept wet via submersion in 200 μL of sterile PBS) until required for use.

**Figure 1.**
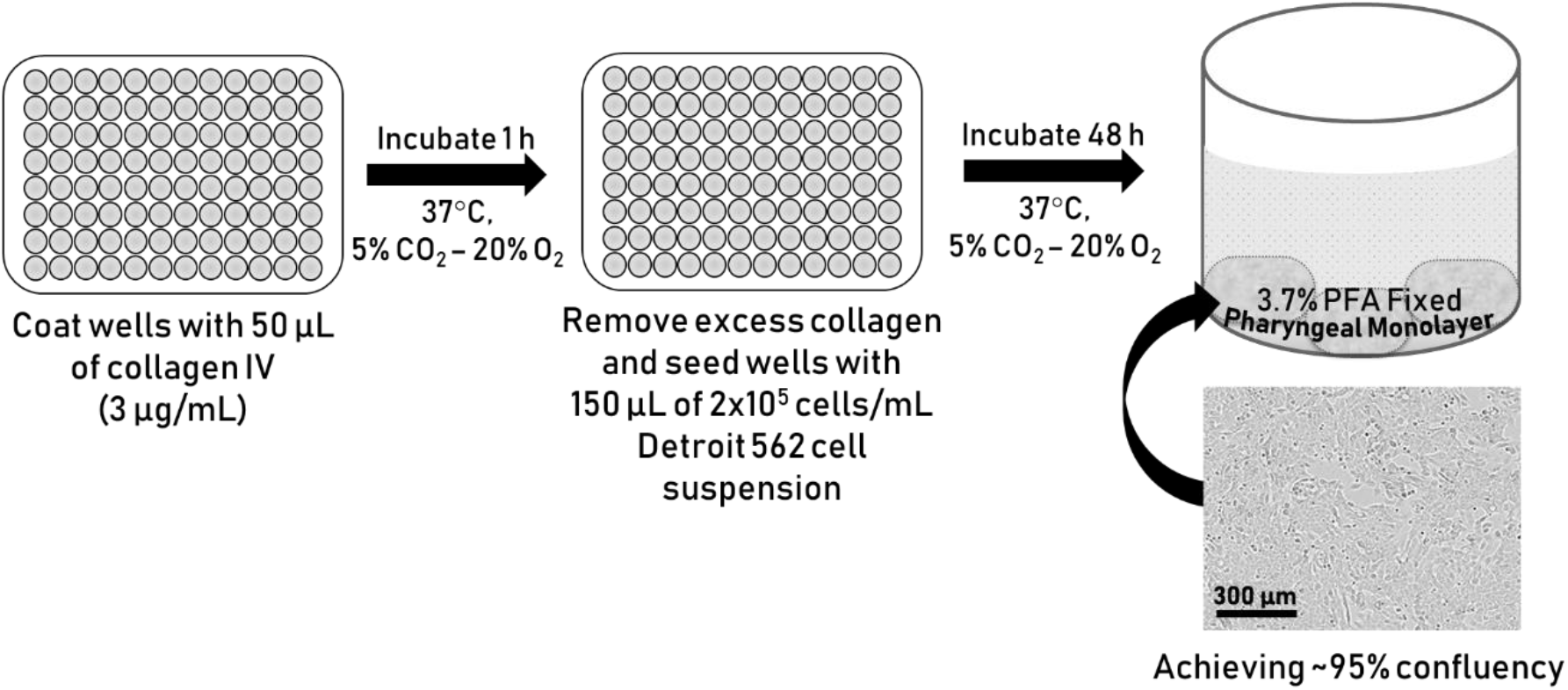
Schematic outlining the process of Detroit 562 pharyngeal cell monolayer formation. Schematic shows collagen coating, seeding with Detroit 562 pharyngeal cells, and finally an example well containing a 3.7% PFA fixed ~95% confluent monolayer of Detroit 562 pharyngeal cells. Example monolayer image taken at 10x objective at the Incucyte.

### GAS biofilm formation

- Wells containing pre-formed fixed Detroit 562 pharyngeal cell monolayers were seeded with 150 μL of overnight GAS culture diluted 1:20 in THY-glucose (0.5% glucose v/v) (THY-G). Wells containing 150 μL sterile THY-G (no bacteria) served as media sterility controls and blanks.
- The plate was incubated for 2 h (34°C, 50 rpm) to facilitate GAS interacting with and adhering to the pharyngeal cell monolayer substratum.

- Tips to avoid evaporation from the biofilm plate:

- Peripheral wells can evaporate quickly, and in turn this affects the biofilms grown and subsequently assayed. Thus, it is advised (where possible) to grow biofilms in the inner-most wells of the plate, with wells not in use filled with 200 μL of sterile water.
- Secondly, store plates in a sealed container filled with water to further avoid dehydration of biofilm wells within the plate.

- Mount on an appropriate stage to keep plate elevated well above water level.
- At 2 h, non-adherent GAS were removed and wells replenished with 150 μL of sterile THY-G. Note: The current protocol was designed to model biofilms formed following the initial interaction occurring between GAS-host cell surface structures at the 2 h point. Thus, this incubation time can be changed dependent on the nature of the study.
- THY-G media was refreshed every 24 h to remove old media containing planktonic/loosely bound cells, debris, and any waste. Note: GAS biofilms are quite delicate.

- Tips for media changing:

- The pipette tip should be placed just below the meniscus of the media, and firmly against the wall of the well at an angle of ~45°.
- To change media with minimal disruption to the biofilm, it is best to remove and replenish the THY-G media slowly and gradually (swapping out 50 μL at a time).

- Together, this reduces any biofilm disruption/dislodgement.
- At 72 h, mature and robust GAS biofilms are produced.

### GAS biofilm biomass crystal violet staining

Biofilm biomass is assayed via crystal violet staining.

- Biofilms were thoroughly air dried for 30-40 mins (or until fully dried)

- Tip: When completely removing media, adjust the pipetting volume (e.g. ~145 μL) to account for any media loss due to evaporation. This prevents/reduces biofilm disruption/loss into pipette tip.
- Dried biofilms were then fixed with 150 μL of 99% methanol for 15 min.

- Tip: Fix with closed lid to minimise methanol evaporation.
- Once fixed, the methanol was gently and slowly removed from the biofilms

- Tip: Given methanol’s propensity to evaporate, pipetting volume adjustment (e.g. ~145 μL) is recommended. This prevents/reduces biofilm disruption/loss into the pipette tip.
- Biofilms were thoroughly air-dried and stained with 150 μL of 0.2% crystal violet (w/v) (Sigma-Aldrich, USA) supplemented with 1.9% ethanol (v/v) for 10 min (RT, static).

- Monolayers with THY (no GAS biofilm) served as media sterility controls and background staining controls, with absorbance values subtracted from those of biofilm samples.
- Once stained, excess crystal violet was removed, and biofilms gently washed twice with 200 μL of PBS.
- Crystal violet stain that had incorporated into the biofilm was re-solubilised upon the addition of 150 μL of 1% sodium dodecyl sulphate (SDS) (w/v) (Sigma-Aldrich, USA).

- Plate was incubated for 10 min (RT, static).
- Biofilm biomass was quantified spectrophotometrically at OD_540nm_ (SpectraMax Plus 384 microplate reader).

### Scanning Electron Microscopy

Scanning electron microscopy (SEM) was utilised to image M1, M12, and M3 GAS biofilms formed on the Detroit 562 pharyngeal cell monolayers as these are all associated with GAS pharyngitis [1]. Preparation of biofilms for SEM was adapted from [20] with the following modifications.

- M1 and M12 GAS biofilms were grown on Detroit 562 pharyngeal cell monolayers previously pre-formed on 13 mm plastic Nunc Thermanox coverslips (Proscitech, USA) in a 12-well polystyrene plate.
- Biofilms were air dried, and pre-fixed in 2.5% glutaraldehyde, 50 mM L-lysine monohydrochloride, and 0.001% ruthenium red solution prepared in 0.1 M HEPES buffer (pH 7.3) (30 min, 4°C).
- Following pre-fixation, biofilms were fixed in fixative solution (2.5% glutaraldehyde and 0.001% ruthenium red prepared in 0.1 M HEPES buffer, pH 7.3) for 1.5 h (4°C) and washed twice in 0.1M HEPES buffer.
- 2% osmium tetroxide vapour was used post-fixation (2 h) followed by three washes with distilled water (each 15 mins).
- A graded ethanol series (30%, 50%, 70%, 90%, and 3x 100%) was then used to remove all water from the biofilms before they were critical point dried (Leica CPD 030, Austria).
- Dried biofilms were then sputter coated with 20 nm platinum (Edwards Vacuum coater, USA) and visualised using a JEOL JSM-7500 microscope (JEOL, Japan) at 500, 5000, and 15 000 x magnification.
- Detroit 562 pharyngeal monolayer controls (without biofilms) were also imaged at 500 and 5000 x magnification.
- Images were taken at random positions within the samples by a UOW Electron Microscopy Centre technician blinded from the study in an effort to reduce bias.

### Statistical Analysis

All statistical analysis was performed using GraphPad Prism (version 8.4.0, GraphPad Software, USA). Datasets were compared using a student T-test, and a P-value of ≤ 0.05 was considered significant.

## Results

### 48 h GAS biofilms are potentiated on Detroit 562 pharyngeal cell monolayers

To exemplify and highlight the need for a more physiologically relevant substratum for GAS biofilm growth, two commonly studied GAS isolates, M1 (5448) and M12 (PRS-8), implicated in GAS pharyngitis were assessed and were the primary focus of this study [1]. Specifically, 48 h M1 and M12 GAS biofilms were assessed for their biofilm forming abilities on both a plastic substratum and Detroit 562 pharyngeal cell monolayers via crystal violet staining (Fig. 2). Both GAS M-types were found to exhibit significantly greater biofilm biomass when grown on the Detroit 562 pharyngeal cell monolayers.

**Figure 2.**
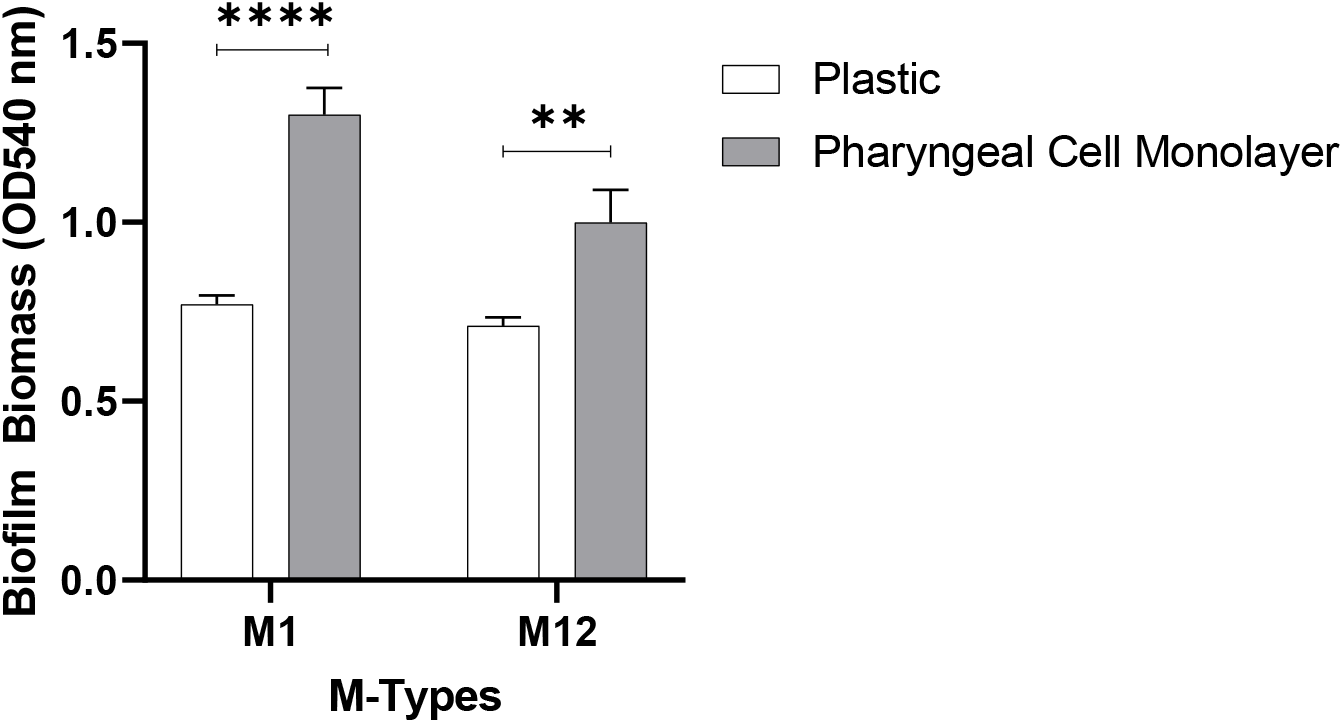
M1 and M12 GAS were assessed for the ability to form biofilm on plastic and Detroit 562 pharyngeal cell monolayers. 48 h biofilms were formed and biofilm biomass ascertained via crystal violet staining. Monolayers with THY (no GAS biofilm) served as media sterility controls and background staining controls, with absorbance values subtracted from those of biofilm samples. Data represents mean ± SEM, ** (P ≤ 0.01) and **** (P ≤ 0.0001); n = 3 biological replicates, with 3 technical replicates each.

### Methanol fixation improves reproducibility of crystal violet staining on GAS biofilms

Crystal violet assays were first described by Christensen, *et al.* [21] as a means of quantifying biofilm. Crystal violet has proven useful in that it detects biofilm in its entirety, staining a biofilms biomass which comprises live and dead cells, as well as EPS matrix. Taken together with its overall ease of use and relatively low cost it has since become a routinely used biofilm stain and detection method [22]. Despite these attributes, there are some drawbacks and limitations to crystal violet use, with concerns around reproducibility [23,24]. Reproducibility can be influenced by a biofilms overall durability and stability, especially during the wash steps [24]. As such, biofilms that are thinner and flimsier require further consideration of overall biofilm handling. Additional steps to the crystal violet assay can be implemented to ensure biofilm retention during the various washing, staining, and de-staining steps of the assay.

In the current study, to minimise disruption and damage to the biofilm, biofilm plate layout/growth conditions and overall handling were optimised as outlined in the methods according to the following specifications; growing biofilms at the inner-most wells of a plate with unused wells filled with water to avoid dehydration; adjusting pipetting volume for media removal to account for dehydration during incubations; placing the plate inside a container containing additional water to reduce media evaporation from the wells; gradual media changes (50 μL at a time, as opposed to the entire 150 μL) etc. Furthermore, to improve the durability of the biofilms during crystal violet assaying, 48 h biofilms grown on the Detroit 562 pharyngeal monolayers were either fixed with methanol or left unfixed and assessed for biofilm biomass (Fig. 3).

**Figure 3.**
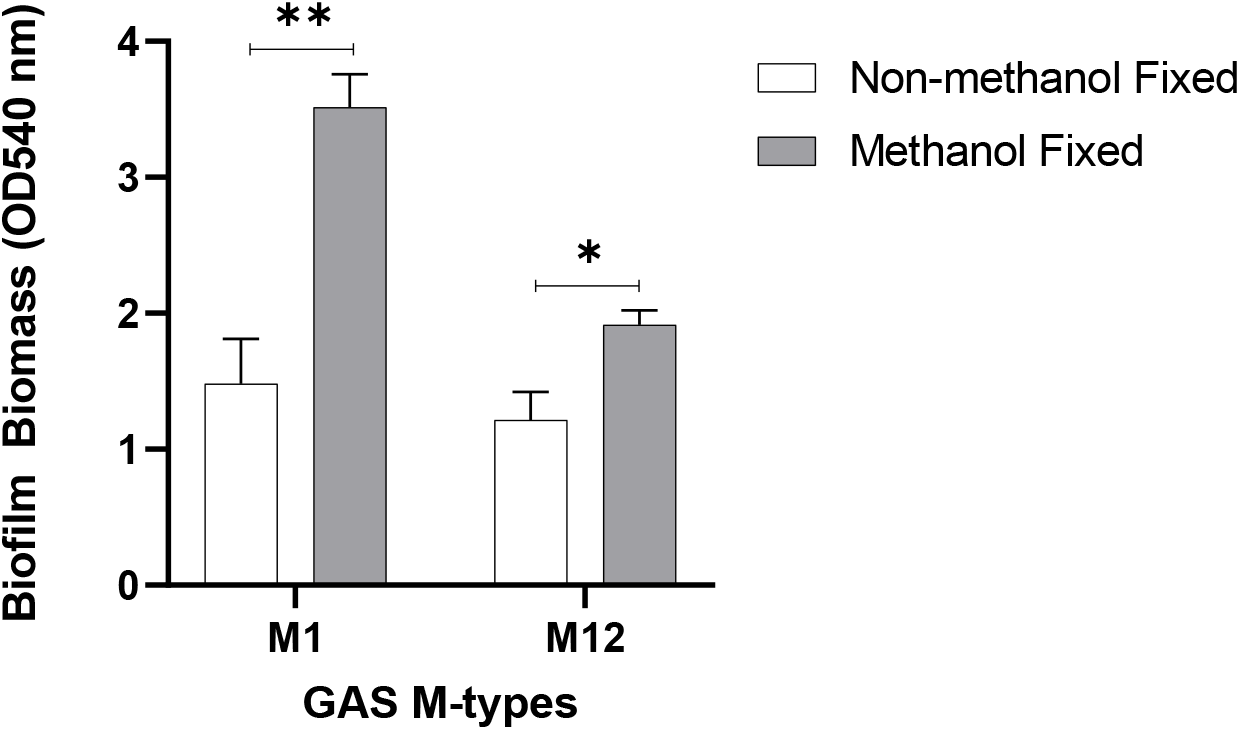
Methanol fixation improves M1 and M12 GAS biofilm biomass detection. 48 h GAS biofilms were formed from planktonic GAS that had initially adhered to the Detroit 562 pharyngeal cell monolayer after 2 h incubation. Biofilm biomass was ascertained via crystal violet staining. Monolayers with THY (no GAS biofilm) served as media sterility controls and background staining controls, with absorbance values subtracted from those of biofilm samples. Data represents mean ± SEM, * (P ≤ 0.05) and ** (P ≤ 0.01); n = 3 biological replicates, with 3 technical replicates each.

Methanol fixation was found to significantly increase retention of biofilm biomass following crystal violet staining (Fig. 3). Hence, for GAS biofilms, we recommend methanol fixation as an additional step prior to crystal violet staining.

### 72 h growth yields optimal biofilm biomass

After optimising the crystal violet assay for GAS biofilms, M1 and M12 GAS biofilm formation on the Detroit 562 pharyngeal cell monolayers was further assessed at extended growth periods of 72 and 96 h to see if greater biofilm biomass was achievable (Fig. 4). Both M1 and M12 formed significantly more biofilm at 72 h compared to 96 h where biofilm biomass seemed to diminish. The biofilm biomass at 72 h was also greater than the biofilm biomass formed previously at 48 h.

**Figure 4.**
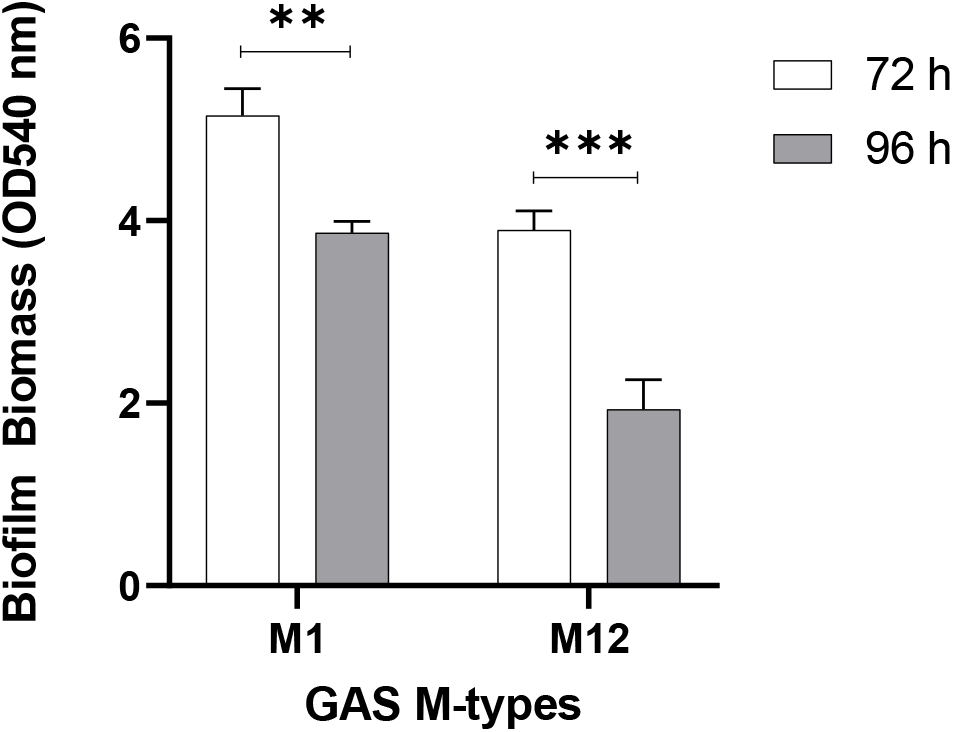
72 h is an optimal period for GAS biofilm formation. M1 and M12 were assessed for GAS biofilm formation at 72 and 96 h. 72 h yielded significantly more biofilm than 96 h. Biofilm biomass was determined via crystal violet staining. Monolayers with THY (no GAS biofilm) served as media sterility controls and background staining controls, with absorbance values subtracted from those of biofilm samples. Data represents mean ± SEM, ** (P ≤ 0.01) and *** (P ≤ 0.001); n = 3 biological replicates, with 3 technical replicates each.

To build upon this and assess the utility of the optimised methodology, additional GAS M types (M3 (90254), M98 (NS88.2), and M108 (NS50.1)) were also assayed for biofilm formation under the same conditions (Fig. 5). As per M1 and M12, both M98 and M108 formed significantly greater biofilm at 72 h compared 96 h. However, biofilm biomass remained relatively unchanged for M3 at both 72 h and 96 h growth periods.

**Figure 5.**
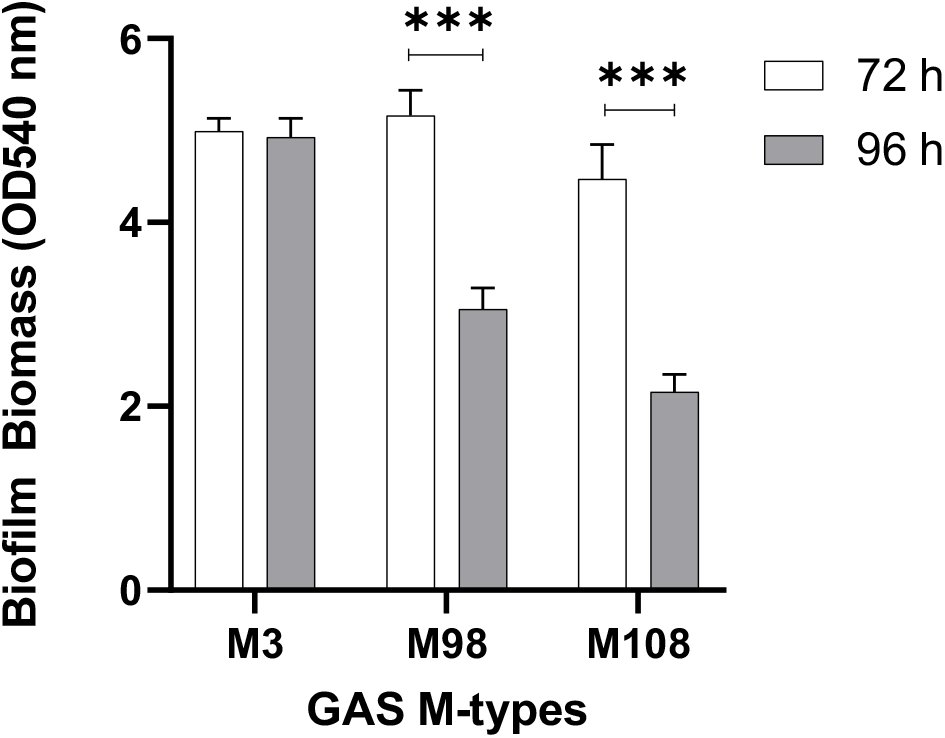
Assessing the utility of the optimised methodology on additional GAS M types (M3, 98, and 108). Biofilm biomass was determined via crystal violet staining. Monolayers with THY (no GAS biofilm) served as media sterility controls and background staining controls, with absorbance values subtracted from those of biofilm samples. Data represents mean ± SEM, ** (P ≤ 0.01) and *** (P ≤ 0.001); n = 3 biological replicates, with 3 technical replicates each.

### SEM imaging reveals 72h M1, M12, and M3 GAS biofilms formed in the host cell-GAS model closely resemble those in vivo

M1 and M12, as well as an M3 strain also implicated in GAS pharyngitis [1] was imaged via SEM. Specifically, 72 h GAS biofilms grown on Detroit 562 pharyngeal cell monolayers were visually observed by SEM for their overall biofilm architecture, arrangement, and structure. M1, M12, and M3 GAS biofilms show cocci chains arranged in three-dimensional aggregated communities atop the Detroit 562 pharyngeal cell monolayers (Fig 6). However, M1 (Fig. 6 A and B) and M3 (Fig. 6 E and H) biofilms were found to arrange in tightly packed aggregates of cocci chains on the Detroit 562 monolayers, whereas M12 biofilms were more loosely arranged atop of the monolayers (Fig. 6 C and D). All biofilms produced noticeable EPS that was found closely associated with the cocci chains (Fig 6. B, D, and F). Detroit pharyngeal cell monolayers(without biofilm)are also shown Fig 6. G and H) depicting the pharyngeal cells arranged in confluent monolayers, with pharyngeal cells displaying their cell surface projections.

**Figure 6.**
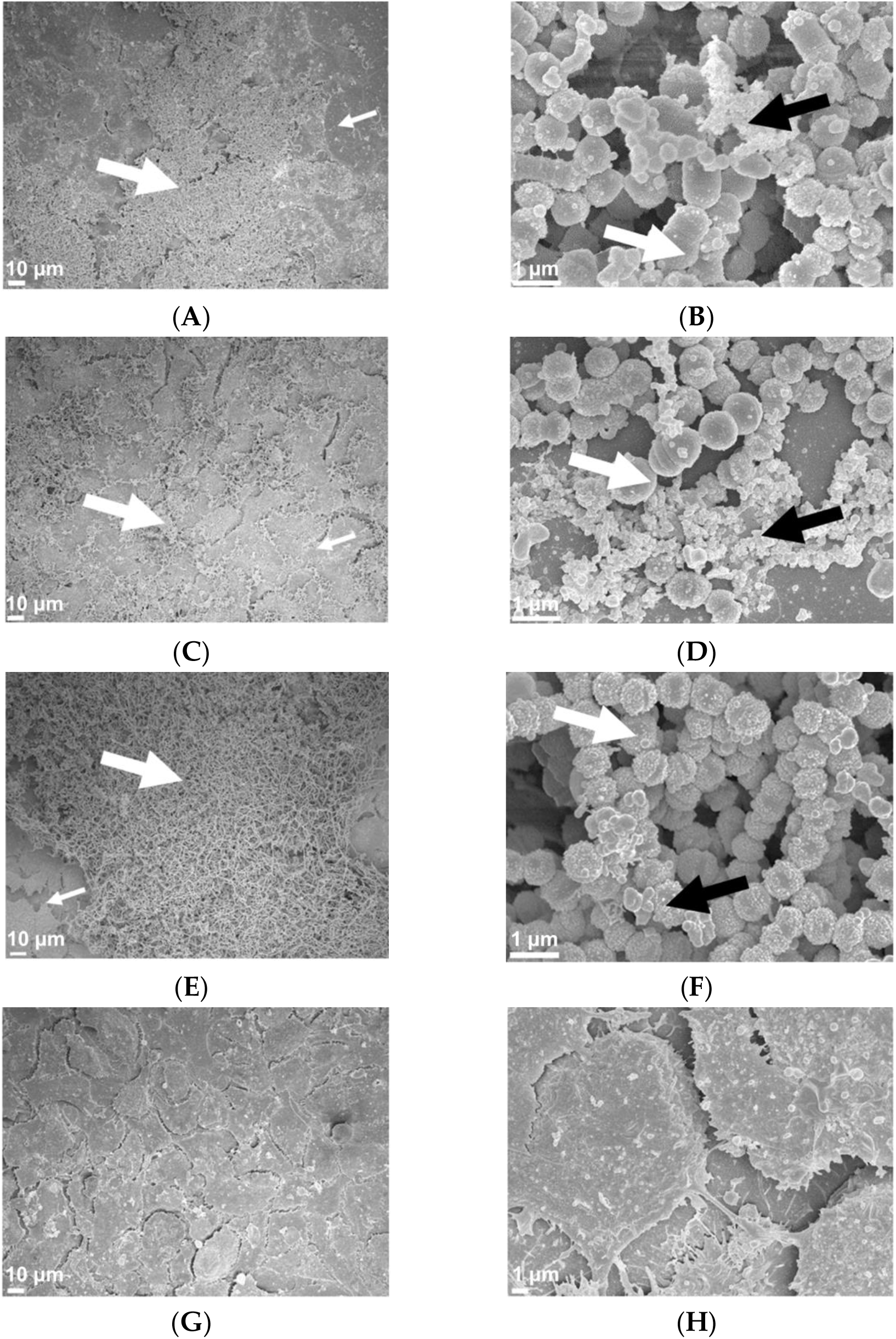
Representative 72 h M1 (A and B), M12 (C and D), and M3 (E and F) GAS biofilms visualised by scanning electron microscopy at 500 and 15 000 × magnification. GAS biofilms show chained cocci (white arrows) arranged into three dimensional aggregated structures with EPS (black arrows) upon the Detroit 562 monolayers (smaller white arrows). Detroit 562 monolayers (without biofilm) (**G** and **H**) were also imaged at 500 and 5000 x magnification. Images represent 3 biological replicates, with 3 technical each.

## Discussion

GAS is a human pathogen, which when in the host is reliant on several host factors to prompt and facilitate dynamic and unique interactions necessary for successful colonisation and persistence. Most *in vitro* plate-based GAS biofilm models used previously do not mimic the host environment. Moreover, the use of an epithelial substratum for growth is rare. Given the importance of the GAS-host tissue interface in mediating the earlier stages of GAS association and adherence, it is likely that these interactions are also crucial for various stages of subsequent biofilm formation and establishment within the host. Thus, modelling in the absence of host factors and/or relevant host epithelial substratum in *in vitro* plate-based biofilm models may result in biofilms that do not accurately represent the GAS biofilms *in vivo*. Here we report an optimised method for GAS biofilm formation using Detroit 562 pharyngeal cell monolayers as a model for GAS-host interaction.

In the current study, an optimised method for forming GAS biofilm on fixed pharyngeal cell monolayers has been developed. Despite Detroit 562 pharyngeal cells not being a primary cell line, they are suitable for this model for numerous reasons; i) the cell line is derived directly from human pharyngeal tissue, a site which GAS readily colonises, ii) retention of surface structures after culturing for adherence (e.g. carbohydrate epitopes) representative of native pharyngeal cells has been noted, and iii) they have been used extensively in planktonic GAS adherence assays [11,25–28]. Overall, utilisation of pharyngeal epithelial substratum within this model aimed to better recreate the host environment than in most of the *in vitro* models used previously.

GAS biofilm biomass increased significantly when grown on fixed Detroit 562 pharyngeal cell monolayers, when compared to biofilm grown on abiotic plastic substratum. This highlights the difference in biofilm potentiation, as a direct result of substratum for biofilm growth. This supports the utility for GAS biofilm modelling to include epithelia as a substratum for biofilm growth over an abiotic plastic surface. Whilst various staining techniques have been developed to assess biofilm biomass, crystal violet staining remains one of the most common techniques within the field. Here, we propose optimisation of this method at the biofilm formation steps preceding crystal violet staining, as well as additional considerations required during the crystal violet assay that are tailored to the GAS biofilms formed using this model. Specifically, to further improve biofilm detection, durability, and subsequent reproducibility of the commonly used biofilm biomass crystal stain assay, methanol fixation was assessed. 48 h GAS biofilms that were methanol fixed prior to crystal violet staining yielded greater biofilm biomass, with less biofilm loss during the intensive staining and washing steps. In turn, reducing error, and increasing reproducibility.

Broth grown GAS is inherently toxic to epithelial cells, as such, there have previously been no models that explore or support long-term GAS biofilm-epithelia co-culture in a plate-based model [15]. Here, we show that fixed pharyngeal cell monolayers support GAS biofilm formation beyond 48 h. However, of the three time points assessed, 72 h is the most optimal biofilm growth period for yielding the greatest biofilm biomass for M1, M12, M98, and M108 GAS. Biofilm biomass was seen to diminish at 96 h, likely resulting from partial disintegration of biofilm. This may be indicative of a mature or older biofilm reaching the final step of the biofilm life cycle - dispersal. Dispersal is thought to be triggered by nutrient exhaustion at the site, which in a host enables bacteria to shift to a motile planktonic state for biofilm re-establishment elsewhere [8,29,30]. These results agreed with a previous study of M6 and M49 biofilms grown on abiotic plastic well surfaces of a 96 well plate-based system, whereby biofilms had greater biomass at 72 h, and exhibited partial disintegration at 96 h with authors attributing this to the age of the biofilm and nutrient limitation [8].

Finally, visual inspection of the GAS biofilms via SEM imaging found M1, M12, and M3 formed biofilm atop the Detroit 562 pharyngeal cell monolayers. Cocci chains, typical of GAS, can be seen arranged in three dimensional aggregated structures coated in EPS matrix for all three M-types. Importantly, biofilms formed in this GAS-pharyngeal epithelial cell model appear similar to SEM images captured of GAS biofilms found at the surface of tonsils removed from patients with recurrent GAS tonsillo-pharyngitis [3].

Here we demonstrate an efficacious GAS biofilm-pharyngeal cell model that can support long-term biofilm formation, with biofilms formed resembling those seen *in vivo*. This model has since been used for assessing the role of pharyngeal cell surface glycans in mediating GAS biofilm formation [12]. Hence, the value of this model is that it can be used to explore a plethora of interactions occurring at the GAS-host cell surface interface and the subsequent effects these interactions exert on biofilm formation.

## Author Contributions statement

Conceptualisation, H.K.N.V, M.L.S-S, and J.D.M; methodology, H.K.N.V; formal analysis, H.K.N.V; investigation, H.K.N.V; resources, M.L.S-S, and J.D.M; data curation, H.K.N.V; writing—original draft preparation, H.K.N.V; writing—review and editing, H.K.N.V, M.L.S-S, and J.D.M; visualisation, H.K.N.V; supervision, M.L.S-S, and J.D.M; funding acquisition, M.L.S-S and J.D.M. All authors have read and agreed to the published version of the manuscript.

## Acknowledgments

The authors acknowledge the valuable assistance of staff at the UOW Electron Microscopy Centre for their help with the specimen preparation and the operation of the JEOL 7500 SEM. We also thank Emma Jayne-Proctor for her technical support.

## References

1. Walker, M.J.; Barnett, T.C.; McArthur, J.D.; Cole, J.N.; Gillen, C.M.; Henningham, A.; Sriprakash, K.S.; Sanderson-Smith, M.L.; Nizet, V. Disease Manifestations and Pathogenic Mechanisms of Group A Streptococcus. Clinical Microbiology Reviews 2014, 27, 264–301, doi:10.1128/CMR.00101-13.

2. Carapetis, J.R.; Steer, A.C.; Mulholland, E.K.; Weber, M. The global burden of group A streptococcal diseases. The Lancet infectious diseases 2005, 5, 685–694.

3. Roberts, A.L.; Connolly, K.L.; Kirse, D.J.; Evans, A.K.; Poehling, K.A.; Peters, T.R.; Reid, S.D. Detection of group A Streptococcus in tonsils from pediatric patients reveals high rate of asymptomatic streptococcal carriage. BMC Pediatrics 2012, 12, 3–3, doi:10.1186/1471-2431-12-3.

4. Akiyama, H.; Morizane, S.; Yamasaki, O.; Oono, T.; Iwatsuki, K. Assessment of Streptococcus pyogenes microcolony formation in infected skin by confocal laser scanning microscopy. Journal of Dermatological Science 2003, 32, 193–199, doi:http://dx.doi.org/10.1016/S0923-1811(03)00096-3.

5. Baldassarri, L.; Creti, R.; Recchia, S.; Imperi, M.; Facinelli, B.; Giovanetti, E.; Pataracchia, M.; Alfarone, G.; Orefici, G. Therapeutic failures of antibiotics used to treat macrolide-susceptible Streptococcus pyogenes infections may be due to biofilm formation. Journal of Clinical Microbiology 2006, 44, 2721–2727.

6. Vyas, H.K.N.; Proctor, E.-J.; McArthur, J.; Gorman, J.; Sanderson-Smith, M. Current Understanding of Group A Streptococcal Biofilms. Current Drug Targets 2019, 20, 982–993, doi:http://dx.doi.org/10.2174/1389450120666190405095712.

7. Oliver-Kozup, H.; Martin, K.H.; Schwegler-Berry, D.; Green, B.J.; Betts, C.; Shinde, A.V.; Van De Water, L.; Lukomski, S. The group A streptococcal collagen-like protein-1, Scl1, mediates biofilm formation by targeting the extra domain A-containing variant of cellular fibronectin expressed in wounded tissue. Mol Microbiol 2013, 87, 672–689, doi:10.1111/mmi.12125.

8. Lembke, C.; Podbielski, A.; Hidalgo-Grass, C.; Jonas, L.; Hanski, E.; Kreikemeyer, B. Characterization of biofilm formation by clinically relevant serotypes of group A streptococci. Applied and environmental microbiology 2006, 72, 2864–2875.

9. Sugareva, V.; Arlt, R.; Fiedler, T.; Riani, C.; Podbielski, A.; Kreikemeyer, B. Serotype- and strain-dependent contribution of the sensor kinase CovS of the CovRS two-component system to Streptococcus pyogenes pathogenesis. BMC Microbiology 2010, 10, 34, doi:10.1186/1471-2180-10-34.

10. Cho, K.H.; Caparon, M.G. Patterns of virulence gene expression differ between biofilm and tissue communities of Streptococcus pyogenes. Mol Microbiol 2005, 57, 1545–1556, doi:10.1111/j.1365-2958.2005.04786.x.

11. Bessen, D.E.; Lizano, S. Tissue tropisms in group A streptococcal infections. Future Microbiol 2010, 5, 623–638, doi:10.2217/fmb.10.28.

12. Vyas, H.K.N.; Indraratna, A.D.; Everest-Dass, A.; Packer, N.H.; De Oliveira, D.M.P.; Ranson, M.; McArthur, J.D.; Sanderson-Smith, M.L. Assessing the Role of Pharyngeal Cell Surface Glycans in Group A Streptococcus Biofilm Formation. Antibiotics 2020, 9, 775, doi:https://doi.org/10.3390/antibiotics9110775.

13. Matysik, A.; Kline, K.A. *Streptococcus pyogenes* capsule promotes microcolony-independent biofilm formation. Journal of Bacteriology 2019, 10.1128/jb.00052-19, JB.00052–00019, doi:10.1128/jb.00052-19.

14. Manetti, A.G.; Zingaretti, C.; Falugi, F.; Capo, S.; Bombaci, M.; Bagnoli, F.; Gambellini, G.; Bensi, G.; Mora, M.; Edwards, A.M., et al. Streptococcus pyogenes pili promote pharyngeal cell adhesion and biofilm formation. Mol Microbiol 2007, 64, 968–983, doi:10.1111/j.1365-2958.2007.05704.x.

15. Marks, L.R.; Mashburn-Warren, L.; Federle, M.J.; Hakansson, A.P. Streptococcus pyogenes biofilm growth in vitro and in vivo and its role in colonization, virulence and genetic exchange. Journal of Infectious Diseases 2014, jiu058.

16. McKay, F.C.; McArthur, J.D.; Sanderson-Smith, M.L.; Gardam, S.; Currie, B.J.; Sriprakash, K.S.; Fagan, P.K.; Towers, R.J.; Batzloff, M.R.; Chhatwal, G.S., et al. Plasminogen binding by group A streptococcal isolates from a region of hyperendemicity for streptococcal skin infection and a high incidence of invasive infection. Infection and immunity 2004, 72, 364–370, doi:10.1128/iai.72.1.364-370.2004.

17. Sanderson-Smith, M.; De Oliveira, D.M.P.; Guglielmini, J.; McMillan, D.J.; Vu, T.; Holien, J.K.; Henningham, A.; Steer, A.C.; Bessen, D.E.; Dale, J.B., et al. A Systematic and Functional Classification of Streptococcus pyogenes That Serves as a New Tool for Molecular Typing and Vaccine Development. The Journal of infectious diseases 2014, 210, 1325–1338, doi:10.1093/infdis/jiu260 %J The Journal of Infectious Diseases.

18. Aziz, R.K.; Pabst, M.J.; Jeng, A.; Kansal, R.; Low, D.E.; Nizet, V.; Kotb, M. Invasive M1T1 group A Streptococcus undergoes a phase-shift in vivo to prevent proteolytic degradation of multiple virulence factors by SpeB. 2004, 51, 123–134, doi:10.1046/j.1365-2958.2003.03797.x.

19. Marks, L.R.; Parameswaran, G.I.; Hakansson, A.P. Pneumococcal interactions with epithelial cells are crucial for optimal biofilm formation and colonization in vitro and in vivo. Infect Immun 2012, 80, 2744–2760, doi:10.1128/iai.00488-12.

20. Williams, D.L.; Bloebaum, R.D. Observing the biofilm matrix of Staphylococcus epidermidis ATCC 35984 grown using the CDC biofilm reactor. Microscopy and Microanalysis 2010, 16, 143–152.

21. Christensen, G.D.; Simpson, W.; Younger, J.; Baddour, L.; Barrett, F.; Melton, D.; Beachey, E. Adherence of coagulase-negative staphylococci to plastic tissue culture plates: a quantitative model for the adherence of staphylococci to medical devices. Journal of clinical microbiology 1985, 22, 996–1006.

22. Wilson, C.; Lukowicz, R.; Merchant, S.; Valquier-Flynn, H.; Caballero, J.; Sandoval, J.; Okuom, M.; Huber, C.; Brooks, T.D.; Wilson, E., et al. Quantitative and Qualitative Assessment Methods for Biofilm Growth: A Mini-review. Res Rev J Eng Technol 2017, 6, http://www.rroij.com/open-access/quantitative-and-qualitative-assessment-methods-for-biofilm-growth-a-minireview-.pdf.

23. Pantanella, F.; Valenti, P.; Natalizi, T.; Passeri, D.; Berlutti, F. Analytical techniques to study microbial biofilm on abiotic surfaces: pros and cons of the main techniques currently in use. Annali di igiene: medicina preventiva e di comunita 2013, 25, 31–42, doi:10.7416/ai.2013.1904.

24. Azeredo, J.; Azevedo, N.F.; Briandet, R.; Cerca, N.; Coenye, T.; Costa, A.R.; Desvaux, M.; Di Bonaventura, G.; Hébraud, M.; Jaglic, Z., et al. Critical review on biofilm methods. Critical Reviews in Microbiology 2017, 43, 313–351, doi:10.1080/1040841X.2016.1208146.

25. Barthelson, R.; Mobasseri, A.; Zopf, D.; Simon, P. Adherence of Streptococcus pneumoniae to respiratory epithelial cells is inhibited by sialylated oligosaccharides. Infection and immunity 1998, 66, 1439–1444.

26. Frick, I.M.; Schmidtchen, A.; Sjobring, U. Interactions between M proteins of Streptococcus pyogenes and glycosaminoglycans promote bacterial adhesion to host cells. European journal of biochemistry 2003, 270, 2303–2311, doi:10.1046/j.1432-1033.2003.03600.x.

27. Ryan, P.A.; Juncosa, B. Group A streptococcal adherence. 2016.

28. Ryan, P.A.; Pancholi, V.; Fischetti, V.A. Group A streptococci bind to mucin and human pharyngeal cells through sialic acid-containing receptors. Infect Immun 2001, 69, 7402–7412, doi:10.1128/iai.69.12.7402-7412.2001.

29. McDougald, D.; Rice, S.A.; Barraud, N.; Steinberg, P.D.; Kjelleberg, S. Should we stay or should we go: mechanisms and ecological consequences for biofilm dispersal. Nature reviews. Microbiology 2012, 10, 39–50, doi:10.1038/nrmicro2695.

30. Gjermansen, M.; Ragas, P.; Sternberg, C.; Molin, S.; Tolker-Nielsen, T. Characterization of starvation-induced dispersion in Pseudomonas putida biofilms. Environmental Microbiology 2005, 7, 894–904, doi:10.1111/j.1462-2920.2005.00775.x.

